# The role of inhibition in resting-state fMRI negative correlations

**DOI:** 10.1101/2024.03.01.583030

**Authors:** Shreyas Harita, Davide Momi, Zheng Wang, Sorenza P. Bastiaens, John D. Griffiths

## Abstract

Resting-state brain activity, as observed via functional magnetic resonance imaging (fMRI), displays non-random fluctuations whose functional connectivity (FC) is commonly parsed into spatial patterns of positive and negative correlations (PCs and NCs). Mapping NC patterns for certain key seed regions has shown considerable promise in recent years as a tool for enhancing neuro-navigated targeting and clinical outcomes of repetitive transcranial magnetic stimulation (rTMS) therapies in psychiatry. These successes bring to the fore several major outstanding questions about the neurophysiological origins of fMRI NCs. In this work, we studied candidate mechanisms for the emergence of fMRI NCs using connectome-based brain network modeling. Simulations of fMRI data under manipulation of inhibitory parameters W_I_ and λ, representing local and network-mediated inhibition respectively, were explored, focusing on the impact of inhibition levels on the emergence of NCs. Despite the considerable difference in time scales between GABAergic neuronal inhibition and fMRI FC, a clear relationship was observed, whereby the greater levels of overall inhibition led to significantly greater magnitude and spatial extent of NCs. We show that this effect is due to a leftward shift in the FC correlation distribution, leading to a reduction in the number of PCs and a concomitant increase in the number of NCs. Relatedly, we observed that those connections available for recruitment as NCs were precisely those with the weakest corresponding structural connectivity. Relative to nominally default values for the models used, greater levels of inhibition also improved, quantitatively and qualitatively, single-subject fits of simulated to empirical FC matrices. Our results provide new insights into how individual variability in anatomical connectivity strengths and neuronal inhibition levels may determine individualized expression of NCs in fMRI data. These, in turn, may offer new directions for optimization and personalization of rTMS therapies and other clinical applications of fMRI NC patterns.

**Author Summary:** Resting-state brain activity, as detected through functional magnetic resonance imaging (fMRI), demonstrates non-random fluctuations in its covariance structure, often characterized as functional connectivity (FC), which is further divided into spatial patterns of positive and negative correlations (NCs). Mapping patterns of NCs of specific key seed regions have demonstrated significant potential as a method for improving the precision of neuro-navigated targeting and enhancing clinical outcomes in the application of repetitive transcranial magnetic stimulation therapies within the field of psychiatry. In our study, we employed the reduced Wong-Wang neural mass model to investigate the physiological underpinnings of NCs observed in resting-state fMRI (rs-fMRI). Our simulated data partially captures the dynamics of empirical rs-fMRI data, revealing that increased inhibition levels correlate with a higher number of NCs. We also observed differential effects on model stability and NCs with varying levels of excitation and inhibition. These findings shed light on the complex interplay between neural dynamics and rs-fMRI connectivity patterns. Importantly, our work contributes to refining model parameters and offers insights for future validation with empirical clinical data. Understanding the factors influencing NCs in rs-fMRI FC has implications for optimizing therapeutic interventions and advancing our understanding of brain function.

## 1. Introduction

### 1.1 Negative correlations in resting-state functional MRI data

The human brain’s spontaneous activity at rest, without any external input, shows non-random, highly structured patterns of organization, which can be measured using resting-state functional magnetic resonance imaging (rs-fMRI). Despite the name, during rs-fMRI recordings, the brain is not ‘at rest’, with neurons and neural populations continuing to exert local and long-range excitatory and inhibitory influences. The most commonly used approach for studying these intrinsic neural fluctuations is by measuring correlations amongst voxel- or region-wise rs-fMRI time series, known as ‘functional connectivity’ (FC). Initially, rs-fMRI FC was reported in the primary motor cortex [1]. Since then, FC patterns have been studied intensely by the neuroimaging community, leading to the progressive discovery that several so-called ‘functional networks’, spanning the entire brain, appear as prominent common variance components in rs-fMRI data. These include the Visual (VN), Somatomotor (SMN), Dorsal Attention (DAN), Ventral Attention (VAN), Limbic (LN), Frontoparietal (FPN), and Default Mode Network (DMN) [2,3]. These networks are consistent across groups of individuals and with different fMRI protocols [4].

In addition to this consistent modular network structure, a prominent feature of rs-fMRI FC patterns is the presence of anti- or negative correlations (NCs), which manifest at both the local (voxel/parcel) level and the regional/network level. It is uncertain what the exact driving source is for these NCs seen in rs-fMRI. Some studies have suggested that the popular preprocessing technique of global-signal regression (GSR) artificially introduces NCs, and therefore their appearance in fMRI data must be interpreted with caution [5,6]. However, other studies have shown strong NCs to be present in rs-fMRI time series both with and without GSR (using a principal component-based noise reduction technique) [7]. Thus, it is likely that at least some of the NCs seen in rs-fMRI FC are driven by real properties of neural dynamics.

A body of evidence suggests that a potential source of NCs in rs-fMRI data could be inhibition, which in turn leads to reduced neuronal activity. For example, in a study looking at responses to electrode stimulation from the visual cortex (V1) of anesthetized monkeys, NCs in the fMRI recordings were linked to decreases in neuronal activity when compared to spontaneous activity beyond the stimulated V1 regions [8]. The authors of this study demonstrated that it is possible to predict the time course of one response based on the other, suggesting a link between them. While this coupling of reduced neuronal activity and NCs does not necessarily imply causation, it does suggest that NCs can potentially be driven by reduced neuronal activity (mediated via inhibition), which in turn leads to reduced local cerebral blood flow. Another comprehensive study employing neuroimaging modalities and electrophysiological recordings highlights this link between the NCs and neuronal inhibition, this time in the somatosensory cortex of anesthetized rats [9]. In this work, it is concluded that neuronal inhibition and concurrent arteriolar vasoconstriction are associated with a decrease in blood oxygenation. This, in turn, could potentially underscore NCs observed in fMRI BOLD data.

### 1.2 Clinical significance of negative correlations

While the exact physiological origins of NCs remain unknown, their clinical importance has been well established, in particular as a tool for improving neuro-navigated targeting in repetitive transcranial magnetic stimulation (rTMS) treatment for major depressive disorder (MDD) [10–12]. Since its introduction in the early 1990s, rTMS protocols for MDD have primarily targeted the dorsolateral prefrontal cortex (dlPFC) with varying degrees of clinical improvements reported [13–16]. In a seminal clinical research study, it was reported that the precise rTMS target location within dlPFC is most effective when it is maximally negatively correlated with the subgenual cingulate cortex (SGC). This was done initially in healthy controls and then tested for reproducibility in an independent MDD patient cohort with MDD [10]. This retrospective finding has subsequently been replicated in prospective clinical trials, across three distinct geographical cohorts, with results staying consistent across different scanner types, preprocessing methodologies, TMS devices, and patient populations [11,12,17].

Despite the clear importance of rs-fMRI NCs for both fundamental and clinical rs-fMRI, there has been relatively little research to date attempting to understand these phenomena in terms of their underlying neurophysiological bases. Related work has studied NCs at a more fine-grained, intra-regional (visual cortex) level, finding that stronger inhibition and weaker excitation can lead to a higher magnitude of NCs in neuronal firing rates [18,19]. Whether these micro-scale NCs bear any direct relationship to rs-fMRI FC NCs, which are significantly slower temporally and larger spatially, is however far from clear.

### 1.3 Present Study

As noted above, previous research has implicated neuronal inhibition generating NCs seen in neuroimaging data [12,18]. We investigated this question using whole-brain connectome-based neural mass modelling of rs-fMRI FC data in the present study. Neural mass models consider the mean activity of an entire neuronal population (representing a brain area or parcel), instead of tens of thousands of individual neurons in that patch of tissue. Connectome-based brain network models combine the activity of several neural masses based on structural (anatomical) connectivity, typically derived from diffusion MRI tractography [20]. This methodology has been used to study resting-state brain activity [21–23], brain oscillations [24], alpha waves [25,26], and traveling waves [20,27], to name a few. Here we developed a connectome-based brain network model employing the reduced Wong-Wang (RWW) neural mass model, as implemented in The Virtual Brain (TVB) software library [28], to study the role of inhibition in generation of rs-fMRI FC NCs. The RWW has been extensively used in prior work to study resting and task-evoked brain dynamics, with the emphasis of modelling studies using RWW typically being rs-fMRI activity (this being what it was designed for; [23,29]). By studying rs-fMRI NCs as a function of inhibition levels in this neural mass model, and focusing on patterns that are consistent across the simulated and empirical rs-fMRI time series, we aim to identify the key mechanistic drivers and organizational principles of this important feature of human brain activity.

In line with prior research, we hypothesize that elevated inhibition levels would lead to an augmented occurrence of NCs in simulated rs-fMRI FC data. Additionally, we explored candidate explanations for the observed spatial patterns of these NCs. Despite the fundamental and clinical significance of rs-fMRI NCs, limited research has delved into comprehending their neurophysiological foundations. A thorough understanding of these phenomena not only holds the potential to enhance clinical applications but also provides insights into the broader dynamics of brain function.

## 2. Results

Using the above-outlined whole-brain computational modelling framework, we explored the role of inhibition, local dynamics, and SC in the generation of negative correlations in the BOLD rs-fMRI time series. In the following, we describe our results using the RWW model. Equivalent simulations were done using the Jansen-Rit (JR) model. This additional analysis was done to validate our results from the RWW model. Our results from the JR model uniformly returned the same conclusions as RWW, which are detailed in the *Supplementary Information* section at the end of this manuscript.

### 2.1 Model simulations fit data well and reproduce the primary characteristics of empirical FC matrices

For each of the 24 subjects, RWW-simulated BOLD FC was generated for a range of values of *G* factor (*Eq. 1 and 4*), and the fit to empirical data (Pearson correlation between simulated and empirical FC matrix upper triangles) was computed. *G* values yielding maximal simulated-empirical FC fits in this brute-force 1-dimensional parameter sweep for each subject were selected for subsequent model runs. This process returned an average fit of R = 0.24 (Std. Dev = 0.06), ranging from R = 0.10 to R = 0.41 (Figure 1A).

**Figure 1:**
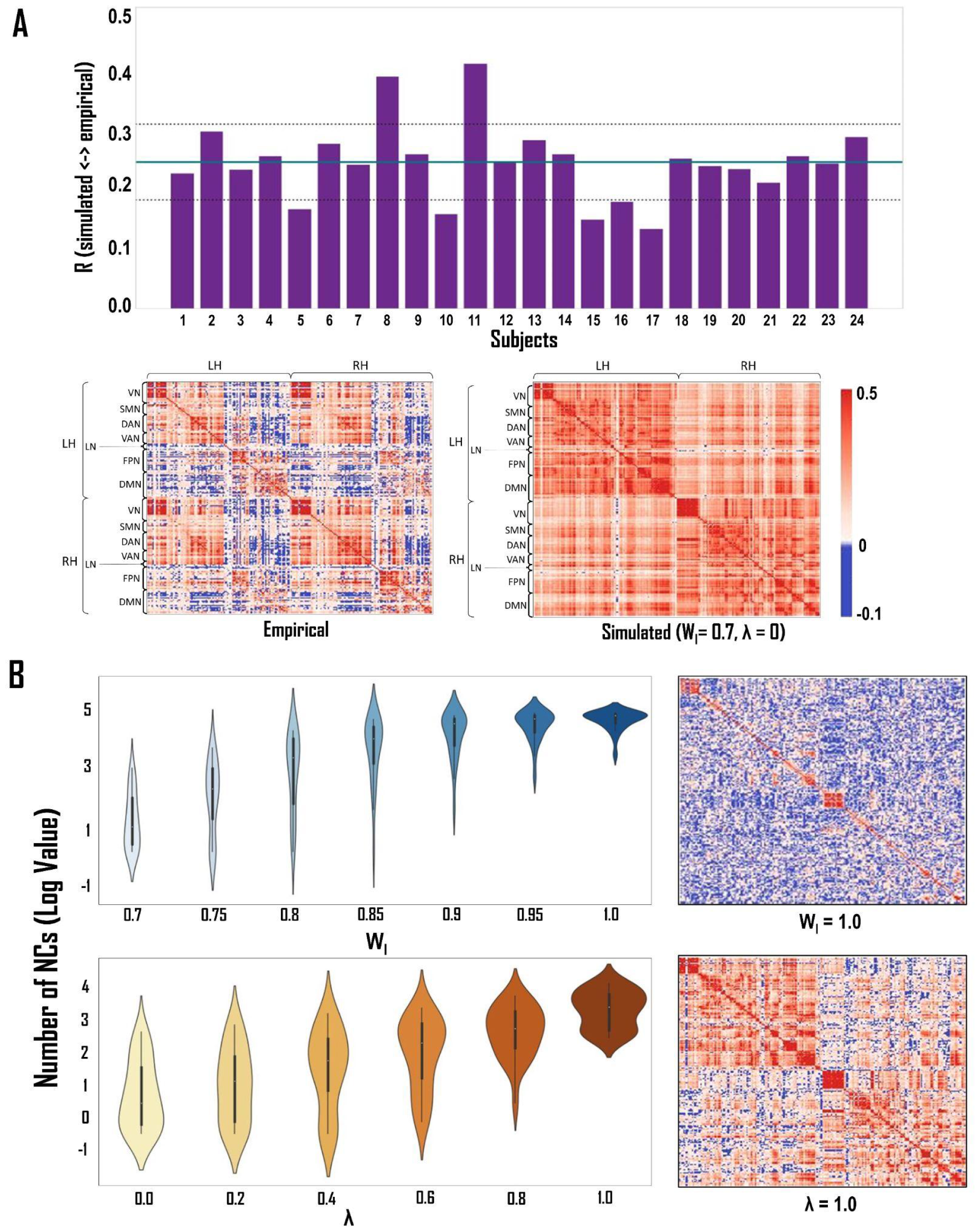
Model Fit to Empirical Data and Inhibition-Induced Changes in Negative Correlations. **A** - *Goodness of fit of the RWW model*. Top: Across our subject cohort, the RWW model showed an average fit of 0.24 ± 0.06 to the empirical data (max. = 0.41). Bottom: The heatmap plots for the empirical and simulated data of a randomly selected subject. The row locations of the canonical Yeo networks are outlined on the left of each FC matrix. **B -** *The effect of increasing the level of inhibition on the number of NCs*. Violin plots show the effect of increasing the level of inhibition on NCs. - *Top:* **W**_**I**_ (external input scaling weight to the inhibitory population); *Bottom:* **λ** (feed-forward inhibition). The y-axis in these plots is represented in the log scale. The heatmap plots on the right showcase another way of visualizing this observation. The increase in the number of NCs is seen when compared to the simulated heatmap in part A. Notably, the FC matrix resulting from elevated feed-forward inhibition levels (λ) on the bottom right shows both qualitatively and quantitatively better fit to the empirical FC matrix on the top left than the optimal FC matrix produced as a result of using the default inhibition values.

The simulated FC from the RWW model can partially capture features of the empirical FC matrix. The model can replicate the FC within some functional networks better than others. For example, the model is better at replicating the FC within the VN when compared to the DMN within the left and right hemispheres. Additionally, the intra-hemispheric simulated FC shows a higher correlation to the intra-hemispheric empirical FC than does the cross-hemispheric simulated FC. Within the left hemisphere, the average fit of the simulated to the empirical data was R = 0.32 (Std. Dev = 0.07), ranging from R = 0.20 to R = 0.48. Within the right hemisphere, the corresponding values were R = 0.30 (Std. Dev = 0.09), ranging from R = 0.10 to R = 0.46. However, the cross-hemispheric fit of the simulated to the empirical was significantly lower than the intra-hemispheric fit. Between the left and right hemispheres, the average fit of the simulated to the empirical data was R = 0.1 (Std. Dev = 0.1), ranging from R = 0.10 to R = 0.38. Hence, the model exhibits a partial replication of the empirical data. When looking at the actual number of NCs, we note that the model does not inherently have many NCs to begin with. The number of NCs increases when the level of inhibition is increased as noted below.

### 2.2 Increasing inhibition increases negative correlations in simulated data

The purpose of selectively manipulating inhibitory parameters in the RWW model was to understand the role inhibition plays in NCs. In the RWW model, we systematically increased the values of W_I_ and λ. Both parameters, when increased, lead to a significant increase in the overall number of NCs. When W_I_ was increased from 0.7 (default value) to 1.0, the mean number of NCs increased significantly (t = -8.19; p<0.0001) from 50.08 ± 92.77 to 13111.25 ± 5313.50. Similarly, when λ was increased from 0 (default) to 1, the mean number of NCs increased significantly (t = -5.35; p<0.0001) from 50.08 ± 92.77 to 2161.00 ± 1859.02 (Figure 1B). The overall fit to empirical data at the maximum inhibition values was significantly reduced to an average of R = 0.18 ± 0.05 (at a W_I_ value of 1.0; t = 3.22; p<0.005; Figure 3B [top]), and remained relatively unchanged at an average of R = 0.24 ± 0.04 at a λ value of 1, across all subjects. Interestingly, in some subjects’ simulated FC matrices, the model fit to empirical data improves slightly following an initial increase in the level of inhibition, but this was eventually seen to decline at higher inhibition levels (Figure 1B (λ heatmap); Figure 3B (top)). This effect was more pronounced following increases in λ compared to increases in W_I_.

**Figure 2:**
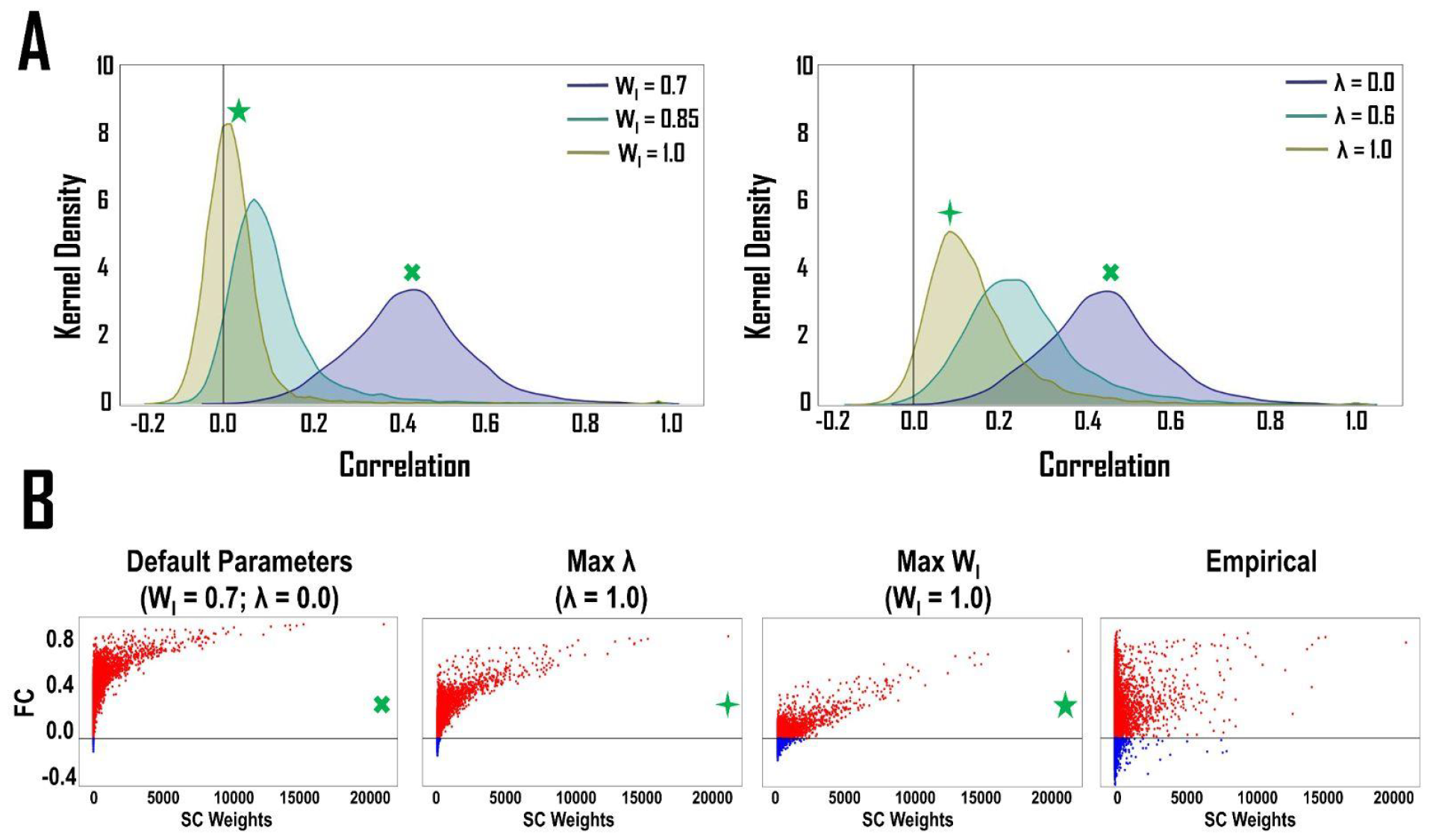
The effect of the level of inhibition on positive and negative correlations. **A** - KDE plots of simulated FC matrix correlation distributions show that increasing inhibition levels leads to a greater number of NCs by reducing the number of PCs. This is seen as an increase in the skew of the right tail of the KDE plot moving from default (navy) through intermediate (teal) to maximum (olive) inhibition parameter values. Furthermore, the KDE plots narrow and shift to the left as a function of the inhibition level. *Left*: Increasing **W**_**I**_ (external input scaling weight to the inhibitory population); *Right*: Increasing **λ** (feed-forward inhibition). **B** - Scatter plots of SC vs. FC matrix values. NCs are seen to have a weaker corresponding SC. As a result, the FC values most likely to become negative at higher inhibition levels, are those with relatively weaker SC. Both plots in this figure represent a single, randomly selected subject; this general pattern was consistently found across the entire subject cohort.

**Figure 3:**
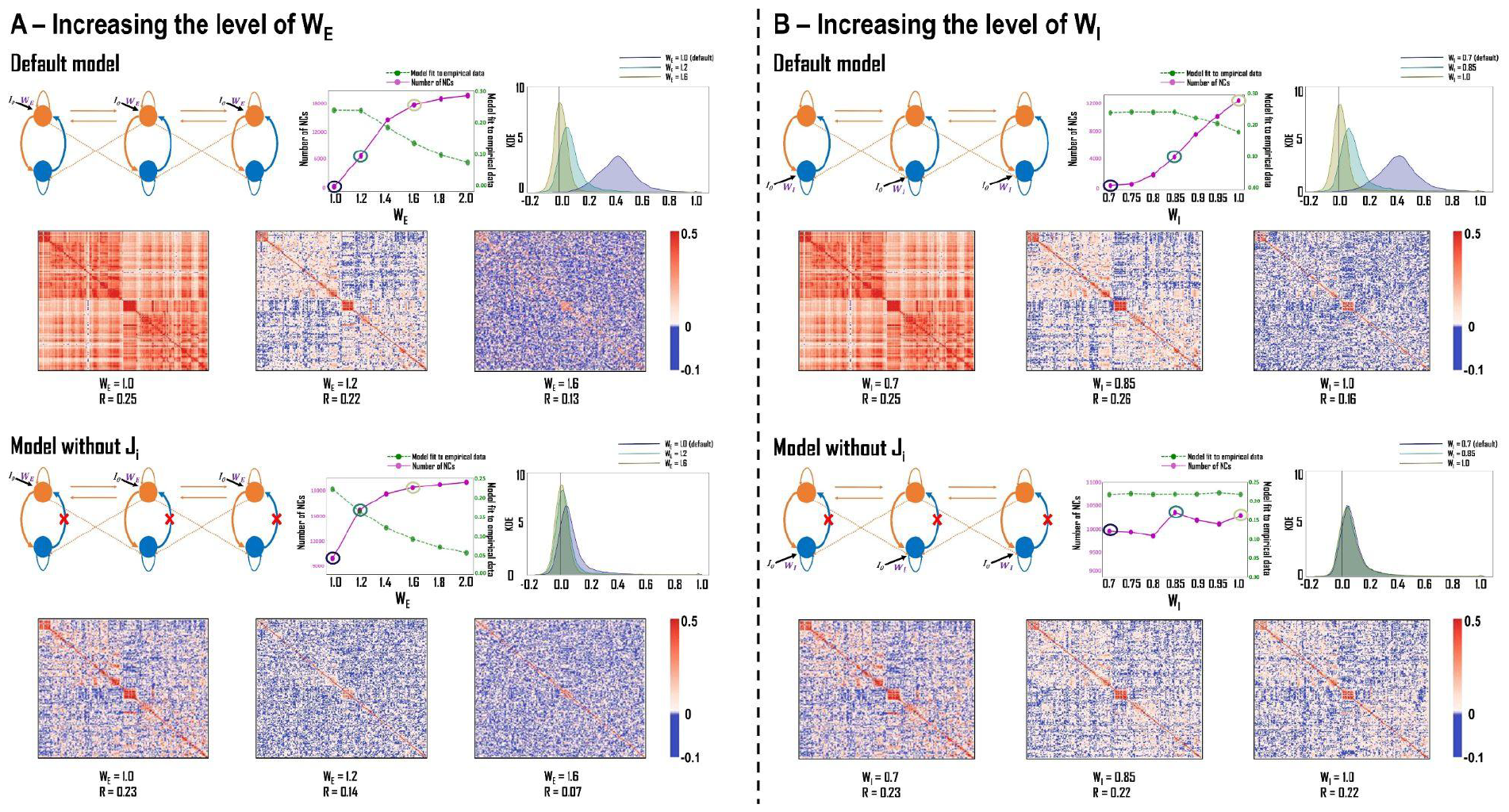
The levels of excitation and inhibition have differential effects on model stability and negative correlations. **A** - *The effect of increasing W*_*E*_ *on model stability and NCs*. ***Top***: When W_E_ (external input scaling weight to the excitatory population) is increased in the default model, there is a concomitant increase in the number of NCs, but this is accompanied by a reduction in model fit (*R*), which for the high W_E_ values leads to random correlations in the simulated FC matrix. ***Bottom***: When J_i_ (inhibitory synaptic coupling) is set to zero and W_E_ is increased, the model starts by being stable and having an increased number of NCs (at the default W_E_ value), but deteriorates at higher W_E_ with the same random structure in the simulated FC matrix seen above. **B** - *The effect of increasing W*_*I*_ *on model stability and NCs*. ***Top***: When W_I_ (external input scaling weight to the inhibitory population) is increased in the default model, initially, there are very few NCs, but there is a significant increase in NCs at higher W_I_ values. While the model stability is reduced, it is not as acute as seen for increases in W_E_ in panel A (top). ***Bottom***: In the model without J_i_, there is a higher number of NCs to begin with (at the default W_I_ value), however, increasing W_I_ does little to affect the overall number of NCs, with only a modest increase at higher W_I_ values. Model fits remain consistent with increases in W_I_.

The increase in NCs was accompanied by a reduction in positive correlations (PCs) in our simulated data when the inhibition parameters were increased. To further investigate this, we studied kernel density estimate (KDE) plots for the overall simulated functional connectivity distribution when inhibition is increased. We observed that increasing inhibition leads to an increase in the skew of the right tail of the distribution. Additionally, there was a narrowing of the distribution and a corresponding shift to the left, at higher inhibition values (Figure 2A). Additionally, when comparing the FC correlations to the SC, we noticed that brain regions showing NCs tend to have fewer direct physical connections between them (i.e., weaker anatomical connections). In other words, stronger SC between brain regions typically leads to PCs between them. As a result, increasing inhibition levels is more likely to cause FC values with lower SC weights to become NCs (Figure 2B).

### 2.3 The levels of excitation and inhibition have differential effects on model stability and negative correlations

The results above demonstrate the effects of increasing inhibitory parameters on NCs. To further understand how inhibition affects simulated data, we conducted control analyses with the RWW model. Control analyses involve modifying an alternative parameter to the inhibitory parameter of interest, to see if variation in NC levels is exclusively determined by inhibitory neuronal processes in the model. We selected W_I_’s excitatory analog, the external input scaling weight to the excitatory population - W_E_.

When W_E_ was increased from 1.0 (default) to 2.0, there was an increase in the number of NCs from 50.08 ± 92.77 to 18359.50 ± 556.62, respectively (Figure 3A [top]). The analysis yielded a significant t-statistic of -146.68, with a p-value of 1.11e-35, indicating a robust result. However, the model’s overall fit to empirical data was significantly reduced from 0.24 ± 0.06 when W_E_ = 1 (default value) to 0.08 ± 0.03 at W_E_ = 2.0 (t-statistic: 3.22; p-value: 1.33e-09). Note that when both W_E_ and W_I_ were increased, there was an increase in the number of NCs. To further understand how W_E_ and W_I_ exert their effects to increase NCs, the excitatory and inhibitory synaptic couplings were manipulated by removing the respective connections (i.e., setting the corresponding synaptic coupling - J_NMDA_ (excitatory) or J_i_ (inhibitory) to 0). In addition to this manipulation, W_E_ and W_I_ were also increased to see how NCs were affected. This led to four different conditions - 1) Increase W_E_, J_NMDA_ = 0; 2) Increase W_I_, J_NMDA_ = 0; 3) Increase W_E_, J_i_ = 0; 4) Increase W_I_, J_i_ = 0. All other parameters in these conditions were set to their respective default values.

Our results from this analysis can be broadly classified into two parts. One where J_NMDA_ was set to 0 and the other where J_i_ was set to 0. When J_NMDA_ was removed, we noticed that the model fit to the empirical data was 0.00. Furthermore, the negative correlations in these simulations were random, not structured, and accounted for approximately half the values in the correlation matrix. This observation was the same when either W_E_ or W_I_ was increased. For W_E_, the mean number of NCs ranged from 19851.50 ± 166.82 at W_E_ = 1.0, to 19866.00 ± 146.48 at W_E_ = 2.0. For W_I_, the mean number of NCs ranged from 19928.50 ± 195.74 at W_I_ = 0.7, to 19929.83 ± 138.33 at W_I_ = 1.0 (*Supplementary Figure 3*). The values reported are averaged values across our 24-subject cohort.

When J_i_ was removed, we observed an increase in the number of negative correlations when W_E_ was increased. When the W_E_ value was increased from 1.0 (default) to 2.0, we observed a significant increase (t = -14.12; p<0.0001) in the number of NCs from 9735.08 ± 3502.60 to 18950.0 ± 422.13, across all subjects. The trade-off with this increase in the number of NCs was a significantly lower model fit to the empirical data. The model fit to the empirical data decreased from 0.22 ± 0.03 at W_E_ = 1.0 (default), to 0.06 ± 0.02 at W_E_ = 2.0, averaged across all subjects (t = 27.04; p<0.0001; Figure 3A [bottom]). When the W_I_ value was increased from 0.7 (default) to 1.0, the number of NCs increased slightly from 9887.58 ± 3324.21 to 9902.08 ± 3356.41. However, this increase was not significant. The model fit to the empirical data was stable and did not change from 0.22 ± 0.03 when the W_I_ value was increased (Figure 3B). Table 1 summarizes the effects of increasing W_E_ and W_I_ on model simulations and the number of NCs for the default model and the model without J_i_.

**Table 1:** The effect of increasing the W_E_ and W_I_ value on model stability and negative correlations. This table complements the information provided in ***Figure 3***. Model stability is represented by FC ✓ (structured correlations) or FC ✗(random correlations). The changes in the number of NCs are compared to the default model, represented by ⇔. An increase in NCs is depicted by ⇑. An excessive increase in NCs (random correlations) is represented with ⇑⇑.

| Parameter | Value | Default Model | Model without $J_i$ |
| --- | --- | --- | --- |
| $W_E$<br>(Scales $I_0$ for excitatory neurons) | Default (1.0) | FC ✓<br>NCs ⇔ | FC ✓<br>NCs ↑ |
|  | Max (2.0) | FC ✗<br>NCs ↑↑ | FC ✗<br>NCs ↑↑ |
| $W_I$<br>(Scales $I_0$ for inhibitory neurons) | Default (0.7) | FC ✓<br>NCs ⇔ | FC ✓<br>NCs ↑ |
|  | Max (1.0) | FC ✓<br>NCs ↑ | FC ✓<br>NCs ↑ |

## 3. Discussion

In this study, we used the RWW brain network model to study the potential mechanistic origins of NCs in rs-fMRI data. Our results point to two cortical microcircuit parameters that influence the number of NCs in simulated fMRI data when varied parametrically. More generally, they provide a proof-of-principle that rs-fMRI NCs can be explored within a whole-brain modelling framework, despite their being an infraslow time series property that is several (temporal) orders of magnitude removed from the underlying neural dynamics. Interesting as these observations are, there are also several important caveats and clarifications in their interpretation, which we discuss below.

### 3.1 The level of inhibition and its impact on negative correlations

By increasing the inhibitory parameter in our model and studying its effects on the model-simulated BOLD data, we aimed to understand the specific role inhibition plays in determining the number of NCs. Interesting observations emerged as we manipulated the levels of inhibition. Firstly, greater levels of inhibition led to a larger overall number of NCs, but this was accompanied by lower overall model goodness-of-fit. The average number of NCs was seen to increase at higher levels of either of the inhibitory parameters - W_I_ (external input scaling weight to the inhibitory population) and λ (feed-forward inhibition parameter). Moreover, varying the levels of specific inhibitory parameters has different effects on the model fit to empirical data. In the case of W_I_, there is a reduction in the model fit to the empirical data across all subjects, but there is a greater increase in the average number of NCs. On the other hand, when the levels of λ are increased, the model fits to the empirical data remains relatively unchanged across our subject cohort, but there is a smaller increase in the average number of NCs. Interestingly, we observed notable patterns in the relationship between inhibitory parameters and the generation of NCs. Examining the empirical data, where the average number of NCs is 3310.83 ± 2404.90 across our subject cohort, we found that the number of NCs at the maximum λ value aligns with this empirical average (2161.00 ± 1859.02), compared to the number of NCs at the maximum W_I_ value significantly exceeds it (13111.25 ± 5313.50). This observation is consistent with our findings that an increase in λ exhibits better fits, more favorable NC patterns, and overall improved model stability compared to an increase in W_I_. An insight into the mechanistic differences between these parameters can be found in *Eq. 4*, highlighting that λ directly influences SC (*C*_*ij*_) between neuronal populations, whereas W_I_ is associated with I_0_, representing neural activity that contributes to the network input. Notably, changes in λ impact network connections in a node-wise manner after input summation, influencing the inputs to each node after summation has occurred. Conversely, W_I_ represents a node-wise input that does not affect the connections between nodes. This distinction likely contributes to the differential effects of W_I_ and λ on the number of NCs, NC patterns, and overall model stability. Thus, the nuanced interplay between these inhibitory parameters underscores their unique roles in shaping resting-state brain dynamics and the emergence of NCs.

Second, our results show that an increase in the number of NCs (due to higher levels of inhibition), is accompanied by a decrease in the number and magnitude of PCs in the simulated data. This is consistent with findings from previous computational modelling research, showing that inhibition is potentially responsible for NCs in electrophysiological studies. For example, increased inhibition and reduced excitation in a spiking network model of the visual cortex led to stronger NCs in the modeled time series [19]. As for the reduction in PCs, this may be occurring as a consequence of enhanced inhibitory feedback, which in turn causes weaker excitatory feedback within the neuronal populations of the model as the amount of inhibition is increased. This amplified inhibitory feedback may thus suppress excitatory neurons, leading to an overall decrease in PCs. Thus, the lower number of PCs observed when NCs are higher may simply result from a lower overall level of interaction and correlation between network nodes. Another explanation for this observation could be that an increase in inhibition can profoundly influence the behavior of neural populations within the brain [29]. This heightened inhibition alters the local dynamics, leading to a reconfiguration of connectivity patterns between neuronal populations. As a consequence, the correlations between these populations are impacted. For instance, stronger inhibitory connections may promote the emergence of NCs, while weakened excitatory connections could reduce PCs. Furthermore, our results show that when inhibition levels are high, FC values with weaker SC are more likely to become negatively correlated (Figure 2B). This suggests that inhibition may play a crucial role in shaping the functional relationships between brain regions, particularly in regions with less direct anatomical connections. These findings underscore the intricate interplay between inhibition, SC, and FC in determining the overall dynamics of the brain network [20].

### 3.2 Excitation and inhibition exert varying effects on model stability and negative correlations

So far, we have seen how the level of inhibition affects the model’s behavior and the number of NCs. To gain a comprehensive understanding of how these effects are mediated, we removed the excitatory and inhibitory synaptic couplings (J_NMDA_ and J_i_, respectively) from our model. Synaptic coupling refers to the interaction or connectivity between different neuronal populations. When considering the RWW model, which consists of a pair of neuronal populations with both excitatory and inhibitory neurons at each of the 200 parcels, synaptic coupling involves the exchange of signals or information through synapses. Excitatory connections (mediated by NMDA in our model) tend to increase the likelihood of an action potential in the target neuron, while inhibitory connections (mediated by GABA in our model) decrease this likelihood. The synaptic coupling between excitatory and inhibitory neurons is a crucial aspect of neural network dynamics. It plays a significant role in regulating the overall activity and balance within the network, influencing factors such as excitability, synchronization, and the emergence of complex patterns of activity. In this study, we deliberately manipulated the synaptic coupling values to investigate the impact of these changes on the overall behavior of the model. By removing the presence of synaptic couplings, and simultaneously increasing the amount of either excitation or inhibition, we sought to elucidate how these factors shape the dynamics, connectivity patterns, and emergent phenomena within the simulated neural network. This approach allows for a comprehensive exploration of how synaptic couplings contribute to the observed behavior and functionality of the model under different conditions.

In Figure 3A, we see how the model behaves when the W_E_ value is increased in two separate scenarios. The first is when the model is unchanged, and the second is when J_i_ is set to 0. When the model is unchanged, increasing W_E_ leads to an increase in the number of NCs. However, in this first scenario, the model’s simulated BOLD fits to empirical data were also significantly poorer at higher W_E_ values. We interpret this result as reflecting a process of runaway excitation, whereby higher values of excitation multiply, causing an imbalance in the model dynamics, loss of synchronous interactions between nodes, and poorer fit to empirical data. In the second case, when Ji was set to 0, and W_E_ was increased, we observed an increase in NCs. However, model stability was quickly seen to deteriorate with increasing W_E_ values. This is because without Ji, there is an uncontrolled activity of excitatory neurons and there is no dampening effect of the inhibitory neurons, unlike the unchanged model. As the W_E_ values are increased, the population activity gets more and more uncorrelated resulting in poorer fits compared to the default model. We also looked at the effect of setting J_NMDA_ to 0 (see *Supplementary Information - Figure 3*). With the link from the excitatory population to the inhibitory population removed, the model becomes a variant of the 1-dimensional excitatory-only system studied in [23], and [30]. In this case, regardless of any increase in W_E_, the model fit to the empirical data was zero, and there was no discernable spatial pattern in the FC matrices, let alone in the NCs. Without J_NMDA_, there is no input from the excitatory neuronal population to the inhibitory ones, and the overall level of excitation within the neuronal population is increased. The resultant decrease in correlation between the excitatory and inhibitory populations [31] leads to a more random, asynchronous activity pattern, as observed in the simulated FC patterns.

In Figure 3B, we see how the model behaves when W_I_ is increased in two analogous scenarios. First, the model is unchanged, and second, when J_i_ is set to 0. When the model is unchanged, increasing the level of W_I_ leads to a significant increase in the number of NCs. However, this increase in the number of NCs is not as high as when W_E_ is increased. However, the model’s simulated BOLD fits to the empirical data remains relatively consistent. When J_i_ was set to zero, and W_I_ was increased, we observed only a slight increase in NCs, but this was nowhere near as potent as the unchanged model. A potential reason for this is that the increase in inhibition levels does nothing to affect the activity of the excitatory neurons as J_i_ is absent. As mentioned above, when J_NMDA_ is set to zero (see *Supplementary Information - Figure 3*), as in the case with W_E_, and for the same reasons [23,30,31], when the level of W_I_ is increased, the model fails to exhibit the expected behavior or ceases to function as intended, with essentially random correlation patterns in the simulated FC matrices. The results from Figure 3 showcase the importance of the interaction between the models’ excitatory and inhibitory populations. Furthermore, these results shed light on the role of the dynamics present in the model and their relationship to NCs.

### 3.3 Implications for the use of negative correlations in clinical neuroscience

In MDD, the predominant and most effective modern neuromodulatory therapies involve rTMS of the left dlPFC and deep brain stimulation (DBS) of the SGC [32]. Previous investigations have revealed that the FC between these targets is characterized by a relatively large number of NCs, suggesting that stimulation of these regions may influence a shared, disease-specific neural network. For example, multiple studies have shown that the NC between the dlPFC and SGC can predict the outcomes of rTMS, with a more negative relationship between these regions leading to a better outcome in patients following rTMS treatment [10,11,33]. Alternatively, brain stimulation has also been shown to induce NCs between brain regions. For example, a study looking at the effects of noninvasive brain stimulation on cortico-thalamic-striatal FC, by applying rTMS to the dorsal-medial prefrontal cortex (dmPFC), showed that MDD patients who showed the most improvement following the treatment course had an increase in NCs between the dmPFC and the insula. These subjects also developed an increased NC between the dmPFC and SGC [34]. In addition, research has demonstrated that various disorders exhibit therapeutic benefits when stimulating sites that manifest NCs (i.e., negatively correlated network nodes). Excitatory stimulation of one site or inhibitory stimulation of the other, both result in symptomatic improvement. Beyond MDD, stimulation of negatively correlated sites has therapeutic effects in Parkinson’s disease, addiction, Alzheimer’s disease, anorexia, gait dysfunction, and pain [32]. Consequently, NCs are significant as they assist in identifying stimulation sites with the potential to induce beneficial effects across various neurological and psychiatric disorders.

Based on the results of this study, the precise picture in terms of how these clinical improvements relate to the neural circuit mechanisms influencing the expression of NCs that we have focused on appears complicated. This is primarily because the topographic distribution of NCs seen in our simulations does not align very well with those in empirical data. While there is an increase in the number of NCs when the level of inhibition is increased, we are not yet able to decipher the specific spatial patterns of NCs or their variations in the population. Future studies can focus on determining the spatial specificity of NCs in model-based simulations and this can in turn shed more light on how NCs contribute to clinical improvements in MDD and other disorders. NCs represent a pervasive characteristic of human brain networks and organization.

They are observed throughout the cortex, irrespective of brain state, highlighting their biological importance. Thus, NCs may hold a particular relevance in clinical contexts.

### 3.4 Limitations

While the results of this study provide some important insights into the potential mechanisms underlying NCs in resting-state brain connectivity, there are some important limitations to highlight.

#### Model Fit to Empirical Data

Concerning model fitting, it is noteworthy that our average model fit to empirical rs-fMRI FC data is 0.24 ± 0.06 (max 0.41) in the RWW model, which is comparatively lower than the previously reported model fits using the RWW model of ∼0.5 [29]. The primary reason for this difference lies in our deliberate choice to employ a higher-resolution brain parcellation in our models, resulting in higher-resolution anatomical and simulated FC matrices. Most studies modeling rs-fMRI with the RWW model have traditionally utilized parcellations with ∼66 brain regions [23,29]. In our work, we opted for the 200-region Schaefer-Yeo parcellation [2,3], which we believe strikes a better balance between spatial granularity and computational feasibility than 66 regions. Our choice of a 200-region brain parcellation in this study, in contrast to the standard whole-brain dynamics studies with fewer regions, introduces a considerable increase in both regions and connections. While a higher ‘R’ value in model fits might suggest a better representation of empirical data, it does not necessarily translate to accurate data prediction [21,35,36]. An unavoidable tradeoff of transitioning to larger parcellations is, however, a reduction in the simulated data fits. The lower fits can be attributed to the following reasons: 1) in the empirical rs-fMRI data, parcel time series are noisier due to averaging over fewer voxels, and 2) in the simulated rs-fMRI data, these additional nodes introduce their own RWW equation noise terms, thereby increasing the total level of stochastic variability in the simulated network. Previous research studies have reported biophysical model fits to empirical data where ∼R = 0.1 to 0.5 [35,36]. In the context of our investigation into the nature of NCs in rs-fMRI data, our focus is not on achieving the best-in-class model fits but rather on obtaining results that are reasonably adequate and qualitatively resemble the empirical data, which our findings demonstrate.

#### Simplified Models

Our study uses the RWW model, which is a simplified mathematical model of brain dynamics. This model does not capture the full complexity and diversity of real brain networks, which display a plethora of features such as oscillations, chaoticity, and spatiotemporal pattern generation. These simplifications and associated assumptions in the models may limit the generalizability of our results to real-world brain functioning. However, we note that we also used the JR model and got similar results (*See Supplementary Information*).

#### Limited Parameters

Our study focuses on the effects of increasing the level of inhibition in the RWW model on the number of NCs in simulated rs-fMRI data. However, other factors and parameters of this model that can potentially affect simulated rs-fMRI data, such as excitation, synaptic connectivity, and network topology, are not fully explored in the study. For example, we did not fully explore continuous variations of J_NMDA_ and J_i_, but rather we set them to zero, as our focus was on the effects of the inhibitory parameters, W_I_ and λ. These and other parameters may interact non-linearly in ways that are significant for the generation of fMRI correlation patterns. Further research on this question should consider a broader range of parameters and their interactions, which may provide a more comprehensive understanding of the physiological origins of NCs.

#### Interpretation of Results

We acknowledge the observed increase in NCs and decrease in PCs with heightened inhibition in the model simulations and we recognize the importance of further investigation to enhance our understanding. The interpretation of the results is grounded in observed correlations within the simulated data and existing literature. To strengthen the reliability of our proposed mechanisms and interpretations, additional research and validation are essential.

### 3.5 Conclusions

This study introduces a novel computational modeling approach to study the effects of varying inhibition and how it affects rs-fMRI dynamics (specifically, NCs) at the population level in connectome-based brain network models. Our work adds to the existing literature on brain network dynamics and provides new insights into the potential mechanisms underlying NCs in brain connectivity. We employ rigorous mathematical modelling techniques, specifically connectome-based whole-brain modelling using the RWW model, to simulate neural dynamics and investigate the relationship between varying levels of inhibition and NCs in resting-state brain connectivity.

Our results show that increasing the amount of inhibition in the RWW model leads to a robust increase in the number of NCs in the simulated data. Different inhibitory parameters in the model had varying effects on the model fit to empirical data and the number of NCs. Previous research has suggested that inhibition could directly or indirectly play a role in determining the number and magnitude of NCs, but it is still unclear why an increase in NCs leads to a reduction in PCs. From a clinical perspective, our findings may have practical relevance and could potentially contribute to the development of novel therapeutic interventions for mental health disorders, such as rTMS therapy for MDD. Potential directions for future research include further validation of the findings using empirical data from clinical studies, exploration of other parameters and factors that may influence resting-state brain connectivity, and clarification of the underlying mechanisms and interactions between inhibitory and excitatory dynamics in the brain.

## 4. Methods

### 4.1 Overview

Resting-state FC is typically used to characterize the spatiotemporal patterns of neural dynamics in the human brain. We use theoretical models to help understand the link between underlying structural connectivity (SC) and FC. The first step in building these models is to determine the density of white matter tracts connecting various brain regions (each represented by a neural mass), depicted by the SC matrix to construct the connectome. This is typically done using diffusion-weighted MRI (dw-MRI) tractography. In the majority of previous work, SC matrices used have had relatively low anatomical resolution of 50-100 brain regions (parcels) [23,29,37–39]. In this study, we use a higher resolution 200 x 200 SC matrix, based on the Schaefer parcellation atlases [3] with dw-MRI data from the Human Connectome Project (HCP) [40]. Figure 4 shows a schematic of our approach. The model equations and neuroimaging data analysis processing details are described below.

**Figure 4:**
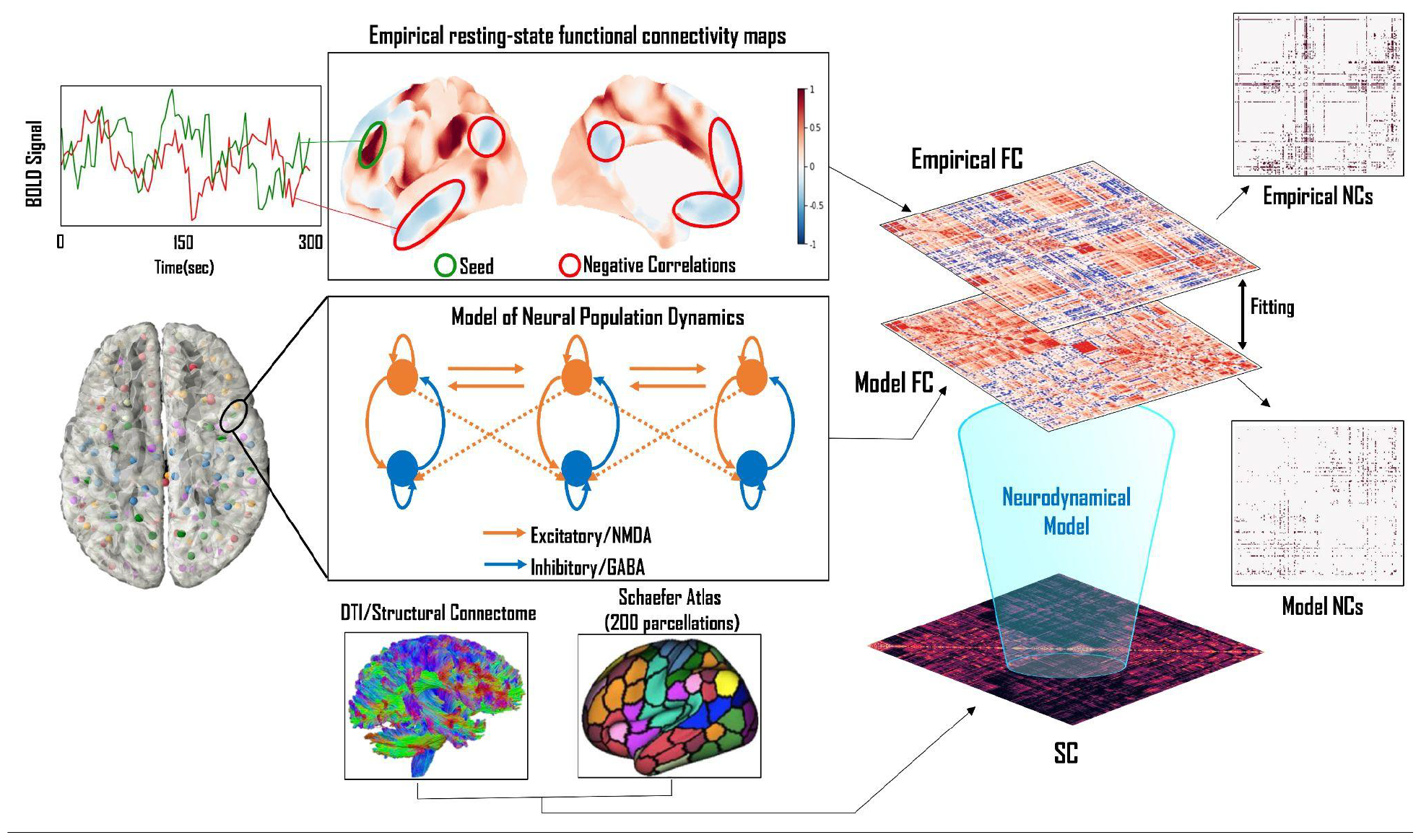
Schematic of the methodological approach. We extend the whole-brain modelling methodology from previous studies to focus on the role of inhibition in NCs in resting-state fMRI data. The RWW model, governed by a set of stochastic differential equations coupled according to the SC matrix, was used to simulate resting-state BOLD fMRI time series data. We first ran a brute-force sweep on the global coupling parameter to place the model in a dynamical regime that approximates empirical fMRI FC well. We then manipulated specific inhibitory parameters in the RWW model to study their effects on the occurrence of NCs in simulated BOLD time series.

### 4.2 Neuroimaging data analyses

DW-MRI data were extracted from 24 randomly selected subjects from the HCP WU-Minn cohort database [40]. The data preprocessing pipeline was run on Ubuntu 18.04 LTS and included tools in FMRIB Software Library (FSL 5.0.3; www.fmrib.ox.ac.uk/fsl; [41]), MRtrix3 (www.MRtrix.readthedocs.io) [42] and FreeSurfer [43]. All DW-MRI data downloaded were already corrected for motion via FSL’s EDDY [44] as part of the HCP minimal preprocessing pipeline [45]. The multi-shell multi-tissue response function [46] was estimated using a constrained spherical deconvolution algorithm [47]. Simultaneously, the T1w images, already coregistered to the b0 volume, were segmented using the FAST algorithm [48]. Following this, anatomically constrained tractography was employed to generate the initial tractogram with 10 million streamlines [49] using second-order integration over fiber orientation distributions [50]. Then, the Spherical-deconvolution Informed Filtering of Tractograms (SIFT2) methodology [51] was applied to provide more biologically accurate measures of fiber connectivity.

Following tractography reconstructions, an anatomical parcellation of the gray/white matter interface was used to segment the whole-brain streamline sets into pairwise bundles, resulting in an anatomical connectivity matrix for each subject. The parcellation used was the 200-node variant of the atlas developed by [3], which incorporates the assignment of each parcel into one of the 7 canonical functional networks identified by [2]- VN, SMN, DAN, VAN, LN, FPN, and DMN. Parcels were defined at the level of surface vertices, mapped to each individual’s FreeSurfer reconstruction using spherical registration [43], and converted to native-space image volumes in register with the dwMRI tractography streamlines. Entries in the resultant 200x200 SC matrices extracted for each subject represent the number of white matter streamlines connecting each pair of ROIs.

### 4.3 Mathematical modeling of neural population dynamics

We modelled activity at each node in the 200x200 network defined by the above tractography analysis using the framework developed by Deco and colleagues [23,29], which employs a mean-field approximation of a leaky-integrate-and-fire spiking network based on the adiabatic assumption that the system is dominated by slow dynamics. As this formulation was first proposed by Wang and colleagues [52,53], we follow others [28] in referring to the resulting equations as the RWW model. The RWW model was first applied to whole-brain fMRI connectivity dynamics by [23], which focused solely on networks of slow excitatory (NMDA) neurons, explicitly omitting fast excitatory (AMPA) and inhibitory (GABA-A) neurons, due to their having significantly shorter decay time constants than NMDA receptors (10 ms vs. 100 ms, respectively). This was then extended in [29] to re-introduce the GABA-A inhibitory neural population, yielding the following two-dimensional system of coupled nonlinear differential equations, representing input currents and population firing rates:

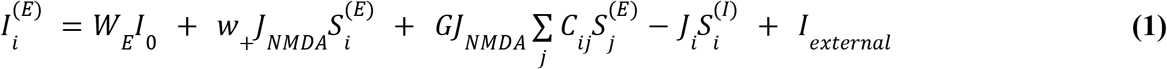

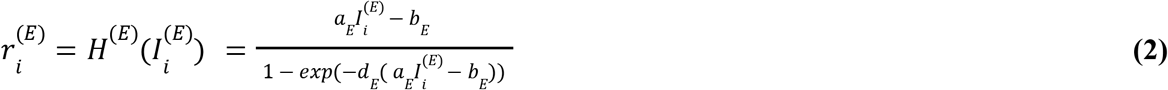

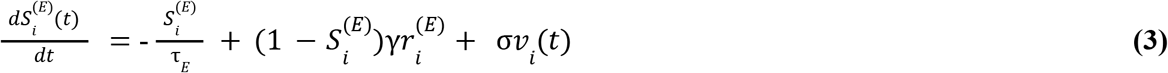

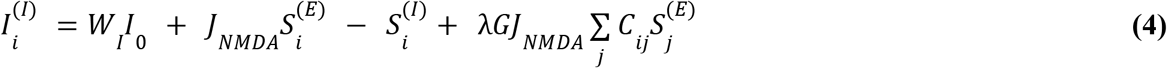

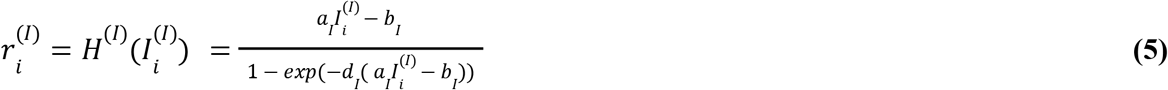

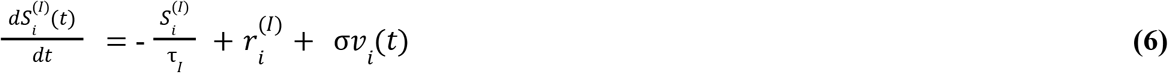

Here, the population firing rate of the excitatory (*E*) and inhibitory (*I*) neuronal populations in a given brain region *i* is given by *r*_*i*_^*(E, I)*^. *S*_*i*_^*(E, I)*^ represents the level of synaptic gating for the *E* and *I* populations, respectively. Synaptic gating refers to the regulatory control or modulation of information flow between neurons at synapses. It involves mechanisms that influence whether a synaptic signal is effectively transmitted or inhibited. These gating processes play a crucial role in shaping neural communication and contribute to the overall dynamics of neuronal networks. *I*_*i*_^*(E, I)*^ are the excitatory/inhibitory input currents to the population *i. Iexternal* is the external stimulation into the ensemble, which is always zero in the present study because we consider only the resting state. A key concept within this model and the present study is that of feedforward inhibition, which refers to the driving of inhibitory population activity by long-range excitatory connections from other nodes in the network. This is represented above by the final terms in *Eq. 4*, and its contribution is scaled parametrically by *λ* ∈ *[0,1]*.

The global scaling factor (G) modulates the strength of inter-areal connections in the neural population model, which are determined by the neuroanatomical matrix (SC), denoted by *C*_*ij*_. The SC matrix specifies the density of fibers between brain regions. *G* offers control over the overall connectivity strength within the global cortical system. In our study, a systematic G sweep explored various values, revealing how *G* influences the dynamics of the model. This exploration sheds light on *G*’s impact on simulated brain dynamics, including synchronization, information flow, and network connectivity. The *G* sweep is crucial for navigating the parameter space and identifying the optimal *G* value that aligns the simulated neural population dynamics with real rs-fMRI data. This iterative parameter optimization enhances the model’s fidelity in replicating empirical observations, providing a more accurate representation of underlying neural processes.

The input-output functions of the excitatory and inhibitory populations are *H*^*(E)*^ and *H*^*(I)*^, respectively. These are mathematical representations of how the activity of neuronal populations responds to incoming inputs. They account for the collective synaptic input received by neurons, incorporating both excitatory *(Eq. 2*) and inhibitory signals (*Eq. 5*). The excitatory (*a*_*E*_, *b*_*E*_, and *d*_*E*_) and inhibitory (*a*_*I*_, *b*_*I*_, and *d*_*I*_) gating variables influence the input currents to the excitatory (*I*_*i*_^*(E)*^) and inhibitory (*I*_*i*_^*(I)*^) populations, respectively. This integrated input is then transformed into a firing rate, which represents the rate at which neurons within the population generate action potentials. The exponential form for the firing rate equation allows for an accurate representation of the nonlinear response profiles of neural tissue, with greater input currents leading to higher firing rates past a certain inflection point. The kinetic parameter is *γ* = 0.641/1000. *I*_*0*_ = 0.382 (nA) is the overall effective external input to the ensemble, which is scaled by W_E_ and W_I_ (the external input scaling weights) for the excitatory and inhibitory populations, respectively. The noise amplitude is *σ* = 0.01 (nA), with *v*_*i*_ being standard Gaussian noise. The local excitatory recurrence is *w*_*+*_ = 1.4. The excitatory synaptic coupling *J*_*NMDA*_ and the local feedback inhibitory synaptic coupling *J*_*i*_ were determined manually using Equations 1-6, to achieve a firing rate of 5 Hz. The corresponding *J*_*NMDA*_ and *J*_*i*_ values were 0.18 and 1.03, respectively. We note that this is slightly above the reported 3.55 Hz but we did not see much difference in the final results [29].

### 4.4 Haemodynamic observation model

The Balloon-Windkessel (BW) model translates simulated neural activity into a simulated BOLD time series, elucidating the hemodynamic processes underlying the neurovascular coupling. The model begins with the neural activity time series as its input, representing the dynamic behavior of neural populations. The hemodynamic response function is a central component, capturing the intricate relationship between changes in neural activity and subsequent alterations in blood flow, volume, and oxygenation within the brain. The “Balloon” component mirrors the compliance of blood vessels, considering the expansion and contraction of vessels in response to cerebral blood flow changes, with parameters such as time constants defining inflow and outflow [54,55]. Simultaneously, the “Windkessel” component represents the damping effect of the vasculature on blood flow, incorporating parameters like arterial transit time [56]. Through a set of differential equations, the model computes the BOLD signal, reflecting fluctuations in blood oxygenation levels across different brain regions.

The signal that induces blood flow is generated by neuronal activity *u*(*t*), where *u*(*t*) is 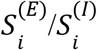 for excitatory and inhibitory neuronal activity, respectively.

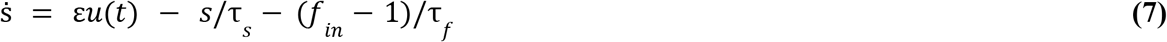

where ε, τ _*s*_, and τ_*f*_ have typical values associated with them.

The inflow of blood *f*_*in*_ is also related to the signal generated by *u*(*t*):

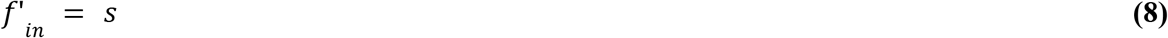

With the inflow of blood, there is extraction of oxygen from the hemoglobin, and the outflow of blood *f*_*out*_ with the change in deoxyhemoglobin (*q*/*q*’).

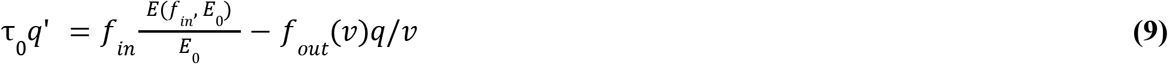

where, *E*(*f*_*in*_, *E*_0_) is the fraction of the oxygen extracted from the inflowing blood, τ_0_ is the time constant, and ν is the volume of venous blood. The rate of change of ν is given by

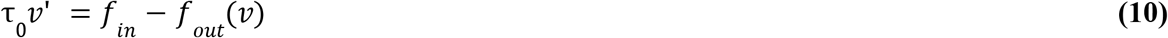

Finally, the resultant BOLD signal *y*(*t*) is a static nonlinear function of normalized ν, normalized total *q*, and the resting net oxygen extraction fraction (*E*0), given by the equation

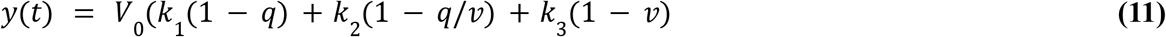

where *k*_1_ = 7 *E*_0_, *k*_2_ =2, *k*_3_=2*E*_0_−0. 2, and *V*_0_ is the resting-blood volume fraction.

To generate a simulated BOLD time series for comparison with empirical data, TVB incorporates an alternative to the differential equations of the BW model described above that is mathematically and computationally simpler without loss of precision. Instead of directly implementing the BW model, TVB utilizes a Volterra Kernel that approximates the behavior of the BW model [57]. The Volterra Kernel serves as an efficient and computationally tractable method to represent the hemodynamic processes involved in converting neural activity to BOLD signals within the context of the RWW model. This methodological choice not only streamlines the simulation process but also maintains the fidelity of the simulated BOLD time series, facilitating a comprehensive comparison with empirical rs-fMRI data.

### 4.5 Exploring the effects of varying the inhibition parameters

Numerical simulation of the RWW equations was carried out using the Virtual Brain (TVB) Python library [58], which is designed to simulate and model large-scale brain networks. Equations 1-6 were used to simulate resting-state BOLD time series within the TVB environment [28]. The focus of this study was to observe the effects of inhibition on NCs in our simulated BOLD time series data. To this effect, except for the control analysis, we exclusively manipulated the inhibitory parameters in the RWW model. The first was the inhibitory input scaling weight - W_I_, which was increased from the default value of 0.7 to a maximum of 1.0 in increments of 0.05. We also increased the feed-forward inhibition parameter - λ from the default value of 0.0 to a maximum of 1.0 in increments of 0.2. All other parameters in the model were left at their respective default values (see above). The inhibitory parameters W_I_ and λ were increased separately to study how increasing each parameter affects the model dynamics and its impact on NCs. In the control analysis for the RWW model, we manipulated W_E_, which is the excitatory analog of W_I_. To study how local dynamics affect NCs, we altered the excitatory and inhibitory couplings - J_NMDA_ and J_i_, respectively. Both the couplings were ‘cut’ i.e., set to a value of zero while we increased either W_E_ or W_I_. This was done to obtain a more comprehensive insight into how these excitatory and inhibitory parameters exert their effects.

## 5. Data and Code Availability

All analyses reported in this paper were conducted on CentOS Linux compute servers running Python 3.7.3, using the standard scientific computing stack and several open-source neuroimaging software tools - principally TVB (simulated BOLD simulations; [58], Nibabel (neuroimaging data I/O; [59] and Nilearn (neuroimaging data visualizations; [60]. All code and analysis results are openly available at *github.com/GriffithsLab/HaritaEtAl2024_tvb-negative-correlations*.

## 6. Author contributions

SH: Conceptualization, Methodology, Formal Analysis, Writing - Original Draft, Writing - Review & Editing, Visualization.

DM: Methodology, Writing - Review & Editing.

ZW: Methodology.

SPB: Writing - Review & Editing.

JDG: Conceptualization, Methodology, Writing - Review & Editing, Supervision, Funding Acquisition.

## 8. Acknowledgments

We are grateful to the Krembil Foundation, CAMH Discovery Fund, and Labatt Family Network for the generous funding support that has made this research possible.

