## Supplemental Information for "The role of inhibition in resting-state fMRI negative correlations"

**9. Supplementary Information**

**9.1 - Replication of RWW model results with the JR model**

***9.1.1 - Jansen-Rit Model Equations***

In addition to using the RWW model, we used the JR model to see how manipulating the level of inhibition affects NCs in the FC matrix generated by this model. While the RWW model has two neuronal populations, the JR model has three - pyramidal neurons, excitatory interneurons, and inhibitory interneurons. Typically, the JR model is used to model electroencephalography (EEG) alpha rhythms and evoked responses [[25,61]](https://paperpile.com/c/WHTumR/XYwKK+lGZZ). While the vast majority of research using the JR model has focused on understanding brain oscillations by modelling EEG activity, this model has also been used to study fMRI and EEG activity concurrently. For example, [[62]](https://paperpile.com/c/WHTumR/UcOZ) proposed a biophysical model that integrated EEG, fMRI, and brain metabolism to understand the relationship between neuronal activity (EEG) and hemodynamic responses (fMRI). In another study, a model-driven approach to studying brain oscillations by integrating EEG and fMRI data used the JR model [[63]](https://paperpile.com/c/WHTumR/bPxI3). In this study, a modified version of the JR model is used to look at FC at the slow (i.e., BOLD/fMRI) timescale. This model is derived from the lumped parameter model [[64]](https://paperpile.com/c/WHTumR/vKDv), where a given cortical column is modelled by three neuronal populations. These are the ‘feedforward’ pyramidal cells, which receive inhibitory and excitatory feedback from local excitatory and inhibitory interneurons, as well as excitatory input from adjacent (or distant) cortical columns. This model details the electrophysiological activity taking place inside a cortical column to study and reproduce EEG alpha activity. The post-synaptic potential of the pyramidal neurons is equivalent to the EEG signal of the model. Two mathematical operators model each neuronal population. The first mathematical operator converts the mean pulse densities of incoming action potentials into an excitatory or inhibitory postsynaptic membrane potential (EPSP or IPSP, respectively). This PSP operator linearly transforms incoming impulses and is given by -

$h_{e}(t)= \left\{ Aat e^{-at} t \geq0 \right\}$	**(7)**

$= \left\{ 0 t \leq0 \right\}$

for excitatory cases, and

$h_{i}(t)= \left\{ Bbt e^{-at} t \geq0 \right\}$	**(8)**

$= \left\{ 0 t \leq0 \right\}$

for inhibitory cases. *A* and *B* are the maximum EPSP and IPSP amplitudes, respectively; while $a$ and $b$ represent the combined effects of the membrane's reciprocal time constant and other distributed delays in the dendritic network. The second mathematical operator takes the mean membrane potential of the neuronal population (i.e., the output from the neuronal population) and converts it into an average pulse density of action potentials. This is given by a nonlinear function, in the form of a sigmoid,

$Sigm(v) = 2e_{0}/[1+e^{r(v_{0}-v)}]$.	**(9)**

The maximum firing rate on the neuronal population is given by e_0_. The PSP at half the maximum firing rate is given by v_0_, and r is the slope of the sigmoid transform [[25]](https://paperpile.com/c/WHTumR/XYwKK).

Four connectivity constants characterize the interaction between the three neuron subtypes in the JR model (i.e., pyramidal cells, excitatory, and inhibitory interneurons). These constants are α_1_, α_2_, α_3_, and α_4_, and they account for the total number of synapses between the interneurons and the axons and dendrites of the cortical column neurons. Including these connectivity constants, the following six differential equations describe the JR model:

$ý_{0}(t) = y_{3}(t)$	**(10)**

$ý_{3}(t) = Aa Sigm[y_{1}(t)-y_{2}(t)] - {2ay}_{3}(t) - a^{2}y_{0}(t)$	**(11)**

$ý_{1}(t) = y_{4}(t)$	**(12)**

$ý_{4}(t) =Aa\{p(t)+\alpha_{2}C*Sigm[\alpha_{1}C*y_{0}(t)]\}-{2ay}_{4}(t)-a^{2}y_{1}(t)$	**(13)**

$ý_{2}(t) = y_{5}(t)$	**(14)**

$ý_{5}(t) =Bb\{\alpha_{4}C*Sigm[\alpha_{3}{C*y}_{0}(t)]\}-{2by}_{5}(t)-b^{2}y_{2}(t)$	**(15)**

where $y_{0}$, $y_{1}$, and $y_{2}$ are the outputs of the PSP block from the pyramidal cells, the excitatory interneurons, and the inhibitory interneurons, and $y_{3}$, $y_{4}$, and $y_{5}$ are their first-order derivatives, respectively. *C* is a connectivity constant that is equal to 135 and is multiplied by a certain factor, given by the alpha values to characterize the connections between the three neuronal populations of the JR model.

The above equations describe the JR model for a single population. We modified this model in our study to account for multiple populations based on our parcellation resolution (i.e., 200 regions based on the Schaefer Atlas). This is based on the equations of the double-column model, which contains two versions of equations 10-15 but can have different system parameters ($A$, $B$, and $v_{0}$) for each population [[25]](https://paperpile.com/c/WHTumR/XYwKK). The modification involves creating a custom TVB model class (c.f. GitHub repository for this study) that extends the standard TVB JR model by adding a PSP block to each node and using its output as a coupling variable. The output is scaled by the SC and then used as an input to equations 10-15. Specifically, this is represented by the following equations:

$ý_{6}(t) = y_{7}(t)$	**(16)**

$ý_{7}(t) = Aa_{d} Sigm[y_{1}(t)-y_{2}(t)] - {2a_{d}y}_{7}(t) - a^{2}y_{6}(t)$	**(17)**

where $a_{d}$ ≈ $a/3$. This represents the delay in the time constant of the input/output signals between two populations.

In the JR model, we manipulated the inhibitory parameters - B and α_4_. We increased B, which is the maximum amplitude of the IPSP from a default value of 22.0 to a maximum of 22.6 in increments of 0.2. We also increased α_4_, from a default value of 0.25 to a maximum of 0.253 in increments of 0.01. This parameter is proportional to the number of synapses made by the inhibitory feedback loop (inhibitory interneurons) to the dendrites of the feedforward neurons (pyramidal cells). In the control analysis for the JR model, we manipulated α_2_, which is the excitatory analog of α_4_, i.e., it is proportional to the number of synapses made by the excitatory feedback loop (excitatory interneurons) to the dendrites of the feedforward neurons (pyramidal cells). We increased α_2_ from a default value of 0.8 to a maximum of 0.803 in increments of 0.01. This was done to see how varying the excitatory parameter affected the NCs in the simulated BOLD of the JR model.

***9.1.2 - Effects of varying inhibition in JR simulations***

Compared to the RWW model the JR model simulated FC showed a lower overall fit to the empirical data, with an average fit of 0.19 ± 0.04, and a maximum fit of 0.28 across our subject cohort. However, the JR model produces a higher number of NCs with default model parameters than the RWW model (Supplementary Figure 1A).

In the JR model, we increased B and α_4_. Similar to the RWW results, both parameters, when increased, lead to an increase in NC. For B, when increased from 22.0 (default value) to 22.6, the mean number of NC significantly increased from 10371.6 ± 4139.94 to 19170.5 ± 1344.68 (t = -13.17; p<0.0001). Similarly, when α_4_ was increased from 0.25 (default) to 0.253, the mean NC increased from 10371.6 ± 4139.94 to 16413.8 ± 3596.28 (Supplementary Figure 1B, 2B) t = -12.69; p<0.0001). The model fit to the empirical data was lowered, at the maximum α_4_ value (0.253), to 0.16 ± 0.03 (t = 4.75; p<0.0001). At the maximum B value (22.6), the fit to the empirical data significantly dropped to 0.10 ± 0.04, with some subjects having no proper fit (i.e., ~0), across all subjects (t = 9.30; p<0.0001).

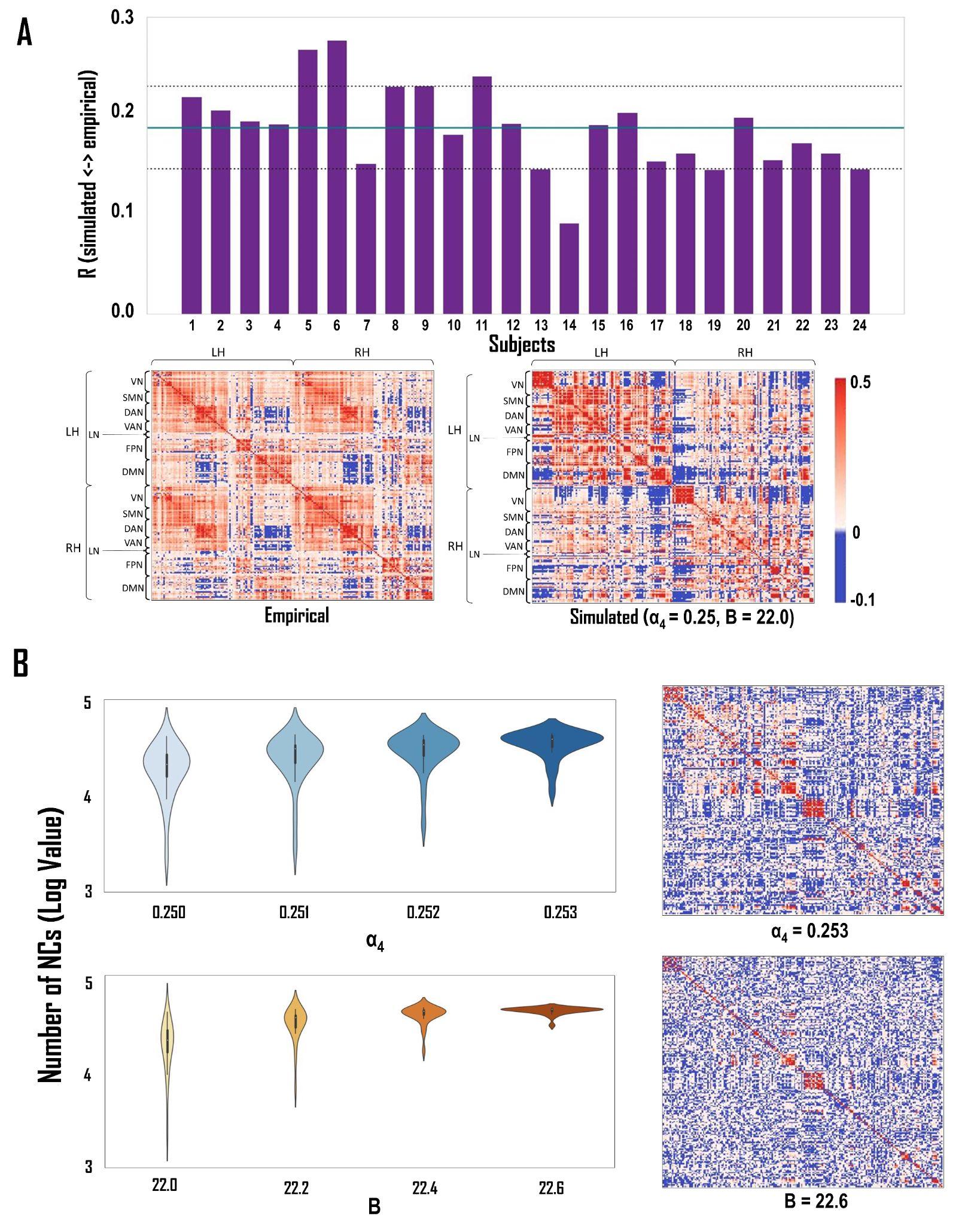

**Supplementary Figure 1: *Model Fit to Empirical Data and Inhibition-Induced Changes in Negative Correlations.* A** - *Goodness of fit of the JR model.* Top: Across our subject cohort, the JR model showed an average fit of 0.19 ± 0.04 to the empirical data (max. = 0.28). Bottom: The heatmap plots for the empirical and simulated data of a randomly selected subject. The row locations of the canonical Yeo networks are outlined on the left of each FC matrix. **B -** *The effect of increasing the level of inhibition on the number of NCs****.*** Violin plots show the effect of increasing the level of inhibition on NCs. - *Top:* **α_4_** (number of synapses made by the inhibitory feedback loop to the dendrites of the feedforward neurons)*; Bottom:* **B** (maximum amplitude of the IPSP). The y-axis in these plots is represented in the log scale. The heatmap plots on the right showcase another way of visualizing this observation. The increase in the number of NCs is seen when compared to the simulated heatmap in part A.

Like the results of the RWW model above, the increase in the number of NCs at greater levels of inhibition also leads to a decrease in the number of PCs. We observed that increasing inhibition leads to an increase in the skew of the right tail of the distribution. Additionally, there was a narrowing of the distribution and a corresponding shift to the left, at higher inhibition values (Supplementary Figure 1A).

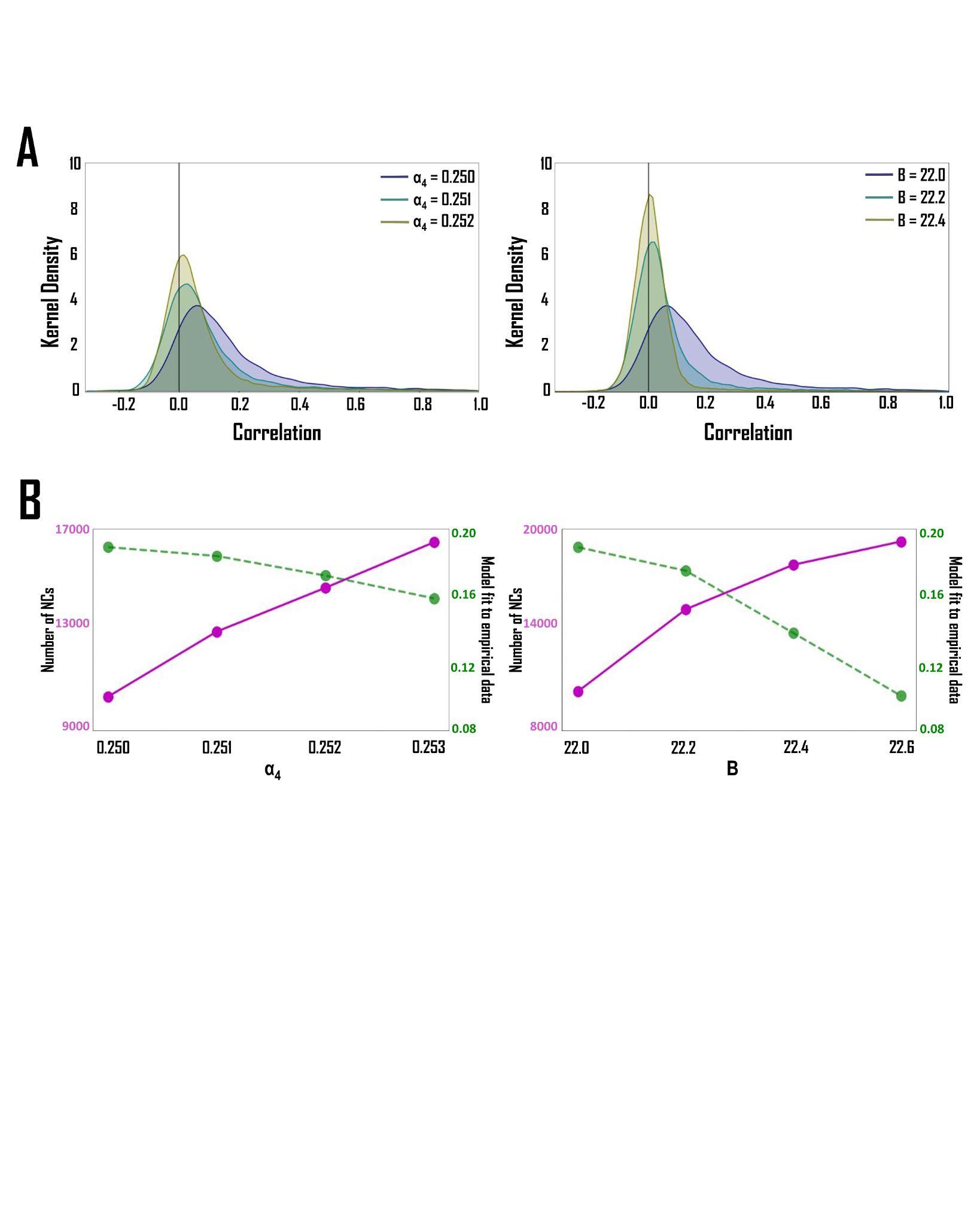

**Supplementary Figure 2: *The effect of the level of inhibition on negative correlations.* A** - KDE plots of simulated FC matrix correlation distributions show that increasing inhibition levels leads to a greater number of NCs by reducing the number of PCs. This is seen as an increase in the skew of the right tail of the KDE plot moving from default (navy) through intermediate (teal) to maximum (olive) inhibition parameter values. Furthermore, the KDE plots narrow and shift to the left as a function of the inhibition level. *Left*: Increasing **α_4_** (number of synapses made by the inhibitory feedback loop to the dendrites of the feedforward neurons); *Right*: Increasing **B** (maximum amplitude of the IPSP). **B** - Line plots depict the increase in the number of NCs as a function of increasing levels of inhibition. The model fit (*R*) to the empirical data is depicted by the dashed green line, and the number of NCs is represented by the pink lines. For both **α_4_** (left) and **B** (right), there is a decrease in *R* with increasing levels of inhibition, but this effect is more pronounced for increases in **B**.

The JR model control analysis involved manipulating α_2_, which is the excitatory analog of α_4_. When increasing α_2_ from 0.8 (default) to 0.803, we saw a slight decrease in NCs from 12081.5 ± 4379.05 to 11350 ± 4289.93. The model fit remained relatively consistent, changing from 0.18 ± 0.05 at α_2_ = 0.8 to 0.17 ± 0.04 at α_2_ = 0.803. These changes in the number of NCs and model fits were not significant. Compared to the results above (when α_4_ was increased), we see that increasing the connectivity of the inhibitory interneurons (α_4_) leads to a significant increase in the number of NCs, and increasing the connectivity of the excitatory interneurons (α_2_) leads to a small decrease in the number of NCs.

***9.1.3 - Comments on the JR model simulation results***

Our simulations with the JR model aimed to investigate the role of inhibition in shaping NCs by adjusting the inhibitory parameter in this model and analyzing its impact on simulated BOLD data. Our observations showed similar trends to the RWW model discussed above, as inhibition levels were manipulated, where higher inhibition levels correlated with a greater overall number of NCs but were also associated with lower model goodness-of-fit. Both B (maximum amplitude of the IPSP) and α_4_ (number of synapses made by the inhibitory feedback loop to the dendrites of the feedforward neurons) showed an increase in the average number of NCs with elevated levels. However, both inhibitory parameters lowered the model fit to empirical data. Increasing α_4_ resulted in reduced model fit across all subjects and a significant increase in NCs. On the other hand, elevating B levels showed an even lower model fit but led to a much higher increase in the average number of NCs across our subject cohort.

Similar to our results obtained from the RWW model, in the JR model, increases in NCs following greater inhibition levels saw a decrease in PCs. The decrease in PCs may be attributed to intensified inhibitory feedback, weakening excitatory interactions within neuronal populations. This enhanced inhibitory feedback may suppress excitatory neurons, resulting in an overall reduction in PCs. Additionally, increased inhibition can alter the behavior of neural populations, leading to a reconfiguration of connectivity patterns and impacting correlations between populations. Specifically, stronger inhibitory connections may promote NCs, while weakened excitatory connections could diminish PCs. These findings highlight the complex interplay between inhibition, SC, and FC in regulating brain network dynamics.

Concerning the effects of increasing the level of α_2_ and α_4_, increasing the connectivity of the inhibitory interneurons (α_4_) increases the average number of NCs. Increasing the connectivity of the excitatory interneurons (α_2_) leads to a slight decrease in the average number of NCs. The model fit to the empirical data is consistent in both cases. The reasons for these findings could be attributed to the fact that the JR model has three neuronal populations (the RWW model has only two). The excitatory and inhibitory parameters increased in the JR model involving the respective interneurons, while the pyramidal cell (which is also excitatory) output parameters were not changed. While there are obvious differences between the RWW and JR models, a similar explanation can be offered for why we see these results. Increasing the connectivity of the excitatory interneurons leads to an increase in the strength of the connections between the excitatory interneurons and the pyramidal neurons, an increase in the overall level of excitation, and a higher degree of coordinated activity between these neurons. This increase in correlated activity could be responsible for the slight decrease in NCs we observe when α_2_ is increased. However, it is important to note that the degree and nature of correlation patterns in the cortical network are highly dependent on the specific details of the circuitry and connectivity within the network [[25]](https://paperpile.com/c/WHTumR/XYwKK). Conversely, increasing the connectivity of the inhibitory interneurons suppresses the activity of the pyramidal neurons. The overall firing rate of the pyramidal neurons is reduced and this lowers the correlation between the neural populations. This is a potential reason we see an increase in NCs when α_4_ is increased [[25,31]](https://paperpile.com/c/WHTumR/XYwKK+qzQk3). Ultimately, while the RWW and JR models are different, we observe a similar pattern in both models when the level of inhibition is increased. In both models, we notice an increase in the overall number of NCs when there is a higher level of inhibition. Based on the results obtained in this study, the level of inhibition in the RWW or JR model affects the correlated activity of the neuronal populations. If the correlated activity is decreased (as noticed when the level of inhibition is increased), there tends to be an increase in the number of NCs.

***9.2 - Additional Figure for RWW Analysis***

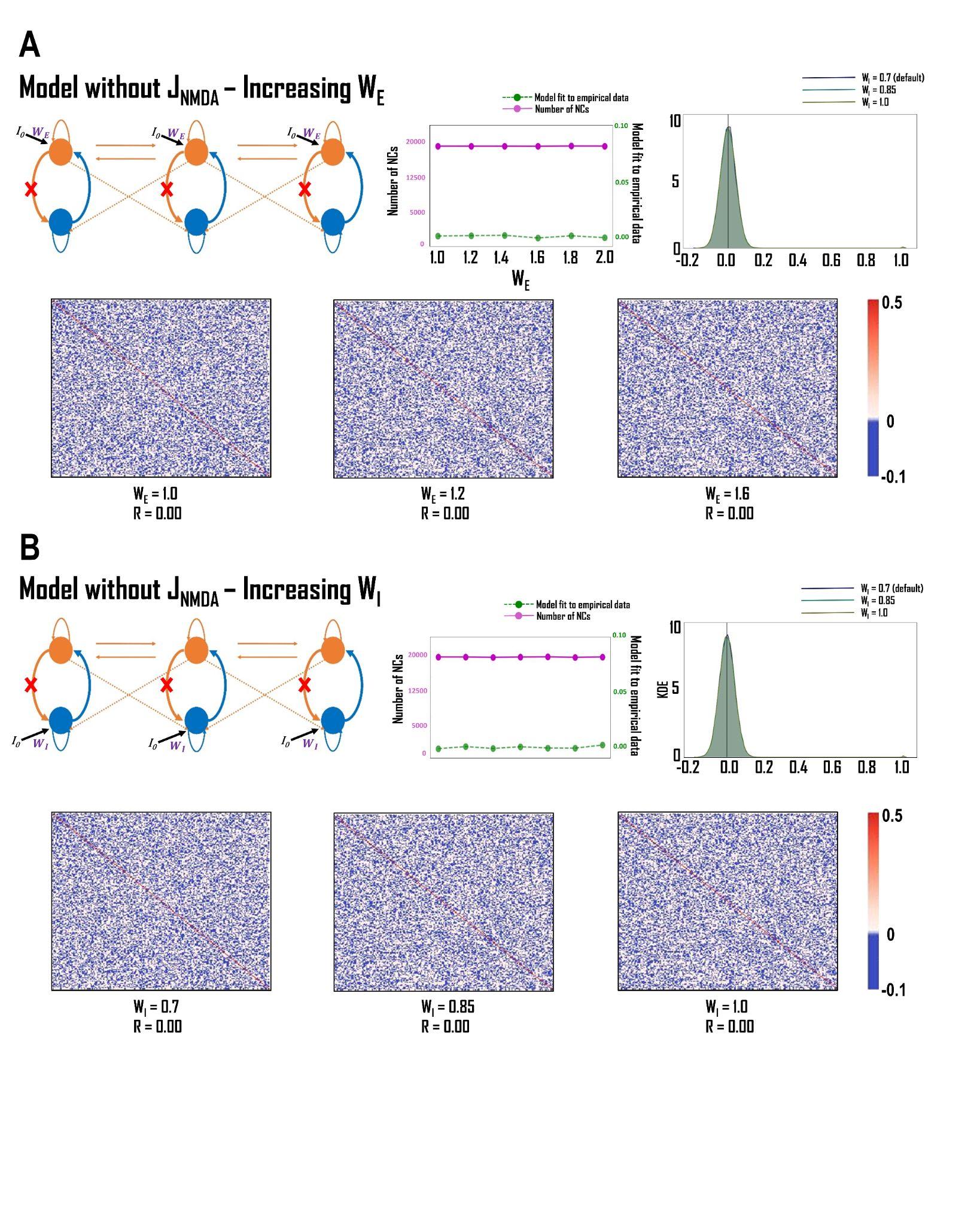

**Supplementary Figure 3: *The effect of removing J_NMDA_ on RWW model simulations.* A** - *The effect of increasing W_E_ on model stability and NCs without J_NMDA_.* When W_E_ (external input scaling weight to the excitatory population) is increased in the model without J_NMDA_ (excitatory synaptic coupling), the model does not behave as intended. The model fit (*R*) to the empirical data is 0.00 and there is no discernable FC pattern. Furthermore, the correlations appear to be random and the distribution of the FC correlations is normal, centered on 0. This is observed from the default through the maximum W_E_ values. **B** - *The effect of increasing W_I_ on model stability and NCs without J_NMDA_.* When W_I_ (external input scaling weight to the inhibitory population) is increased in the model without J_NMDA_, the observations are identical to those described in part A above. The model fit to the empirical data is 0.00, the correlation structure is random, and the FC correlations are normally distributed and centered at 0. This is the case from the default through the maximum W_I_ values.
